# The dynseq genome browser track enables visualization of context-specific, dynamic DNA sequence features at single nucleotide resolution

**DOI:** 10.1101/2022.05.26.493621

**Authors:** Surag Nair, Arjun Barrett, Daofeng Li, Brian J Raney, Brian T Lee, Peter Kerpedjiev, Vivekanandan Ramalingam, Anusri Pampari, Fritz Lekschas, Ting Wang, Maximilian Haeussler, Anshul Kundaje

**Author notes:** equal contribution (Surag Nair, Arjun Barrett, Daofeng Li).

## Abstract

We introduce the dynseq genome browser track, which displays DNA nucleotide characters scaled by user-specified, base-resolution scores provided in the BigWig file format. The dynseq track enables visualization of context-specific, informative genomic sequence features. We demonstrate its utility in three popular genome browsers for interpreting cis-regulatory sequence syntax and regulatory variant interpretation by visualizing nucleotide importance scores derived from machine learning models of regulatory DNA trained on protein-DNA binding and chromatin accessibility experiments.

## Main

High-throughput experimental platforms have revolutionized the ability to profile diverse biochemical and functional properties of biological sequences such as DNA, RNA, and proteins. Nonetheless, the path from data collection to gleaning novel biological insights and generating viable hypotheses is not straightforward. Genome browsers are a crucial tool in this process. By collating multiple data modalities with customizable tracks rendered using intuitive visualizations, genome browsers enable an interactive and interpretable exploration of diverse types of genome profiling experiments and derived annotations. For example, tracks encoded in the BigWig file format are commonly used to visualize continuous profiles such as coverage tracks from high-throughput sequencing experiments like RNA-seq and ATAC-seq, as well as computationally-derived continuous annotations such as sequence conservation. Tracks encoded in the BED file format are frequently employed to visualize sets of discrete genomic intervals of interest such as regions with significant enrichment of experimentally measured signals. More recently, genome browsers have added support for long-range interactions and 2-dimensional views to visualize contact maps from assays such as Hi-C.

However, existing genome browser tracks are not well-suited for intuitive visualization and analysis of DNA sequence features such as transcription factor (TF) motifs. A typical analysis of cis-regulatory elements involves visualizing motif hits, obtained by thresholding scores from position weight matrix (PWM) scanning, as BED-based annotation tracks. While the BED track visualization highlights the genomic coordinates overlapping these motif hits, the specific sequences of each motif hit cannot be gleaned from this track. Instead, one would require cross-referencing a genome sequence track with the motif BED track to identify the sequences of motif hits. Even so, information about motif affinity and if and where the underlying sequence differs from the consensus motif sequence are not immediately apparent. This makes interpretation slow and challenging. This problem is compounded when analyzing multiple cellular states and/or molecular readouts (e.g., ATAC-seq, ChIP-seq) simultaneously, since BED coordinates of enriched regions (peaks) and the set of active TF motifs vary across cell states. Similarly, sequence conservation tracks in non-coding regions often highlight conserved TF binding, but this information is not visible when these tracks are displayed using the standard BigWig rendering. An alternate approach of visualizing DNA bases scaled by heights proportional to meaningful scores such as per-base binding strength has been proposed previously (Schneider, 1997). Such a track can be used to visualize not just the signal score at each base but also the corresponding identity of the bases, thereby highlighting high scoring sequence features at single base-resolution.

Recently, machine learning models that can map DNA sequence to functional readouts from various high-throughput assays have been developed to study the sequence basis of molecular activity and decipher putative functional genetic variants influencing protein-DNA binding, chromatin state, splicing, gene expression, and long-range chromatin contacts (Eraslan *et al*., 2019; Z. Avsec *et al*., 2021; de Almeida *et al*., 2022). These models can be interrogated using various feature attribution methods to infer quantitative, predictive importance scores of each base in any candidate sequence of interest, thereby allowing discovery of predictive nucleotides and sequence features such as TF motifs, splice sites, and polyadenylation sites (Jaganathan *et al*., 2019; Bogard *et al*., 2019). These importance scores are currently visualized in ad-hoc ways that are not suited to seamless exploration and easy sharing.

Hence, we introduce the dynamic sequence (dynseq) genome browser track that displays DNA nucleotide characters at a genomic locus with heights scaled by user-specified, base-resolution, quantitative scores. The dynseq track makes it straightforward to visually recognize sequence features such as TF motifs which are activated in a context-specific manner. The dynseq track is a generalization of previously proposed “sequence walkers” (Schneider, 1997) and is adapted to modern genome browsers. Here, we describe the dynseq track specifications and showcase typical use cases for interpreting cis-regulatory sequence syntax and non-coding, regulatory genetic variants.

To visualize context-specific dynamic importance scores, we first implemented and integrated the dynseq track in the WashU Epigenome Browser (Li *et al*., 2019, 2022). The input file format is the BigWig format (Kent *et al*., 2010) with per base-pair importance scores within broader regions of interest, such as peaks. At each position, the nucleotide character (A/C/G/T) is rendered. Each nucleotide has a distinct color. The height of the character is scaled by the importance score at that position. Negative scores are handled by flipping the character along the x-axis. Hovering the mouse over the track shows the score at that position. Right-clicking on the track opens a customization panel that allows choosing an automatic y-axis scale or setting fixed maximum and minimum limits. When the track is zoomed out such that individual bases cannot be discerned, the track visualization automatically switches to a regular BigWig view. The simple specification makes the dynseq track amenable to be easily incorporated into other genome browsers. We have so far added equivalent native functionality for UCSC Genome Browser (Kent *et al*., 2002) and HiGlass (Kerpedjiev *et al*., 2018) (**Supplementary Table 1**).

To illustrate a typical use case, we trained separate BPNet neural networks (Ž.Avsec *et al*., 2021) to map DNA sequence to base-resolution profiles of DNase-seq and 5 TF ChIP-seq experiments in the K562 cell line (ENCODE Project Consortium, 2012; Davis *et al*., 2018). We used the DeepLIFT feature attribution algorithm to derive nucleotide importance scores for sequences underlying regions with enriched signal (peaks) from each of the models (Shrikumar *et al*., 2017; Lundberg and Lee, 2017). We used the WashU Epigenome browser to visualize and interpret an enhancer in the β-globin locus ~10kb upstream of the HBE1 gene (Kellis *et al*., 2014). We used BigWig tracks to display the observed and model-predicted profiles for each assay, and dynseq tracks to visualize assay-specific importance scores derived from each model (**Figure 1, Supplementary Figure 1**). The dynseq tracks clearly highlight the predicted quantitative influence of different sequence motifs on chromatin accessibility and binding profiles of the 5 TFs. NFE2, GATA1, GATA2 and USF1 binding profiles are predicted to be strongly influenced by their respective direct binding motifs. In contrast, TAL1 binding profile is strongly affected by the GATA motif and multiple weaker TAL1 motifs, which suggests that the GATA TFs are potentially involved in cooperatively recruiting TAL1 to its binding sites. The DNase-seq model’s importance score track highlights the GATA and NFE motifs, suggesting that these motifs, but not TAL and USF, likely drive DNase I hypersensitivity at this locus. We also visualized PhyloP conservation scores using both the standard BigWig rendering and the dynseq track. The PhyloP dynseq track highlights strong conservation of the GATA, NFE, and multiple TAL motifs but not the USF motif. Together, the dynseq tracks are able to reveal subtle, quantitative, context-specific sequence determinants of TF binding and chromatin accessibility thereby providing insights into the architecture of cis-regulatory elements.

**Figure 1.**
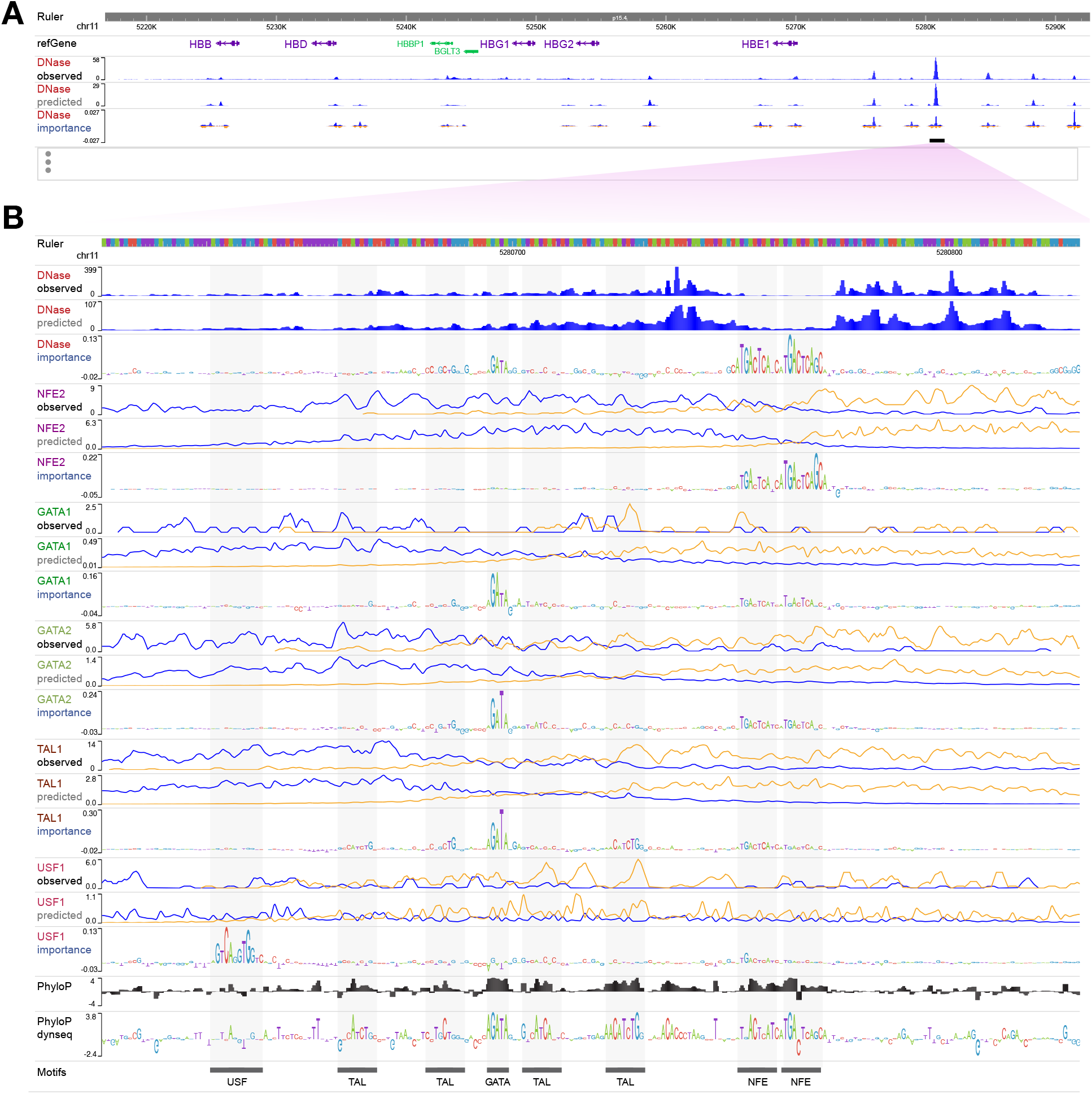
WashU Epigenome Browser session for deciphering sequence architecture of a cis-regulatory element. **(A)** Human β-globin locus with the observed DNase-seq track and zoomed out DNase-seq model-derived importance dynseq track **(B)** An enhancer 10kb upstream of HBE1 (hg38 chr11:5280613-5280820) with observed base-resolution 5’-end coverage tracks of DNase-seq and ChIP-seq targeting transcription factors NFE2, GATA1, GATA2, TAL1 and USF1 in the K562 cell-line, corresponding predicted tracks from BPNet sequence models trained on these data, BPNet model-derived nucleotide importance scores visualized using dynseq tracks, and PhyloP conservation scores visualized using standard BigWig and dynseq tracks. For ChIP-seq observed and predicted tracks, blue denotes the plus (+) strand and orange denotes the minus (−) strand.

Another salient application for dynseq tracks is to explore the sequence features impacted by functional genetic variants. As a case study, we trained BPNet models on ChIP-seq profiles of the SPI1 transcription factor and ATAC-seq profiles in the GM12878 cell line (ENCODE Project Consortium, 2012; Davis *et al*., 2018). We used these models to predict the allelic effect of non-coding variants on ATAC-seq and SPI1 ChIP-seq profiles and derive nucleotide-resolution DeepLIFT importance scores for the sequences containing the reference and alternate alleles of each variant. rs5764238 is a single nucleotide variant that has been previously shown to have a significant allelic effect on SPI1 binding in a binding quantitative trait locus (bQTL) study (Tehranchi *et al*., 2016). We used the Resgen HiGlass browser to visualize the predicted SPI1

ChIP-seq and ATAC-seq profiles as BigWig tracks and the DeepLIFT importance scores as dynseq tracks for the pair of sequences containing the reference and alternate allele of this variant **(Figure 2)**. The BigWig tracks show that the alternate (G) allele is predicted to enhance SPI1 binding and chromatin accessibility, in agreement with the bQTL study. The dynseq tracks show that the G allele creates a strong SPI1 binding motif with high importance scores from the SPI1 ChIP-seq and ATAC-seq models, thereby revealing this motif as the primary driver of enhanced signal. Hence, the dynseq tracks add an additional level of interpretability to the study of genetic variants by focusing attention on visually recognizable sequence features in a context-specific manner.

**Figure 2.**
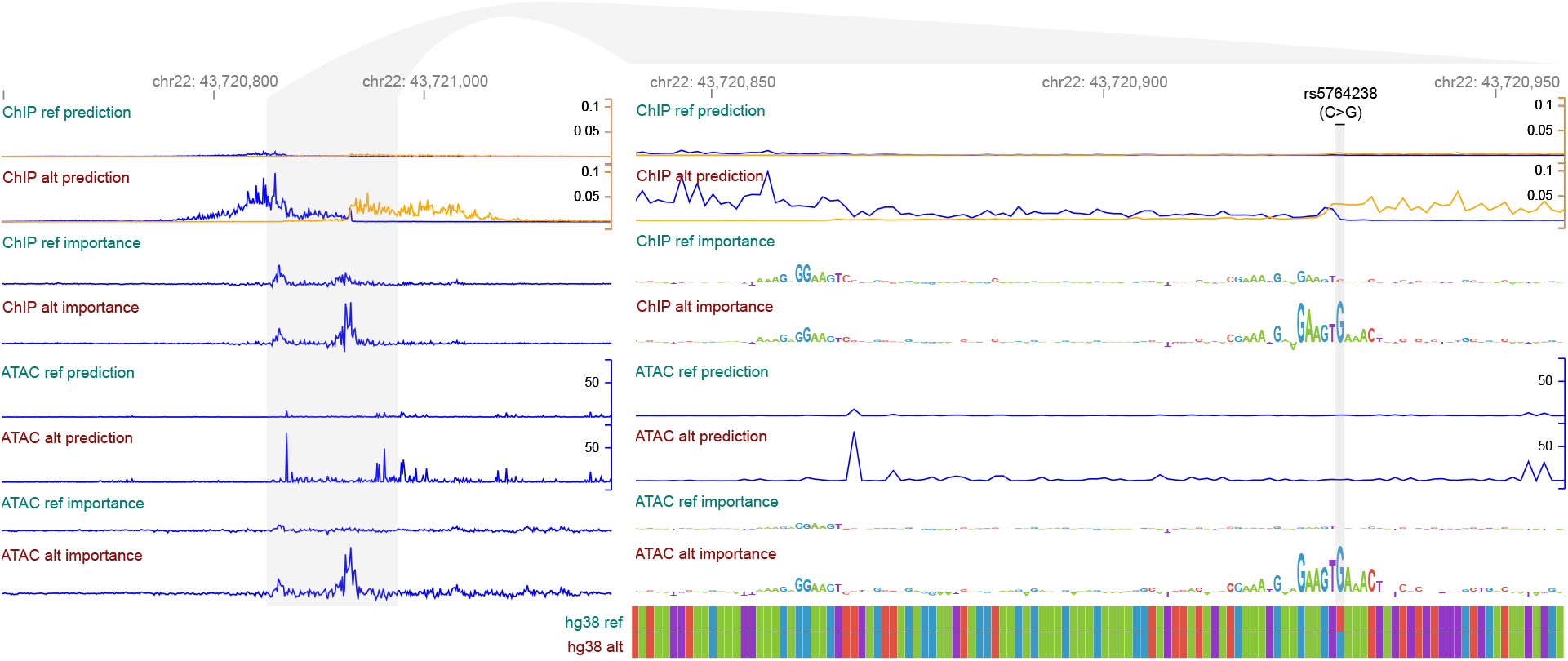
Resgen/HiGlass browser session for model-guided, non-coding variant interpretation. Zoomed out (left, hg38 chr22:43720597-43721177) and close-up (right, chr22:43720840-43720958) views of predicted base-resolution coverage profiles (as BigWig tracks) and nucleotide importance scores (as dynseq tracks) from BPNet models of SPI1 ChIP-seq and ATAC-seq data for a genomic sequence containing the reference (C) and alternate (G) allele of the rs5764238 genetic variant, previously identified as a SPI1 binding QTL. For ChIP-seq predicted tracks, blue tracks denote the plus (+) strand coverage and orange tracks denote the minus (−) strand coverage. The G allele is predicted to increase SPI1 ChIP-seq and ATAC-seq signal by creating a strong SPI1 motif, as seen in the dynseq tracks.

Genome browsers are an indispensable tool for the analysis of genomic and biochemical profiling experiments. We expect that the dynseq tracks will enhance exploratory analysis, discovery and hypothesis generation using genome browsers by enabling contextual interpretation of informative sequence features in genomic elements and those disrupted by genetic variation at single nucleotide resolution.

## Availability and implementation

The dynseq track takes input files in the BigWig format. A tutorial on how to use the dynseq tracks on various browsers is available at https://kundajelab.github.io/dynseq-pages/. The dynseq track is currently available for the following browsers:

- **UCSC Genome Browser** (https://genome.ucsc.edu): Usage information is available at https://genome.ucsc.edu/goldenPath/help/BigWig.html#dynseq.
- **HiGlass** (https://higlass.io) and **Resgen** (https://resgen.io): Dynseq is implemented as a plugin track for these browsers. Source code and usage information is available at https://github.com/kundajelab/higlass-dynseq/.
- **WashU Epigenome Browser** (https://epigenomegateway.wustl.edu): Source code for the WashU Epigenome Browser with dynseq track support is available on GitHub (https://github.com/lidaof/eg-react). Track usage is described in the WashU Epigenome Browser documentation (https://eg.readthedocs.io/en/latest/tracks.html#dynseq).

See **Supplementary Table 1** for more information on functionality supported by each browser.

The data and models used to create the vignettes are available at https://doi.org/10.5281/zenodo.6582100. Code is available at https://github.com/kundajelab/dynseq-paper.

## Acknowledgements

This work was supported by National Institutes of Health grant numbers U01HG009431, U01HG012069 to A.K.; R01HG007175, U01CA200060, U24ES026699, U01HG009391, UM1HG011585, U41HG010972, U24HG012070 to T.W.; 5U41HG002371 to B.J.R., B.T.L., M.H.

---

**Supplementary Figure 1.**
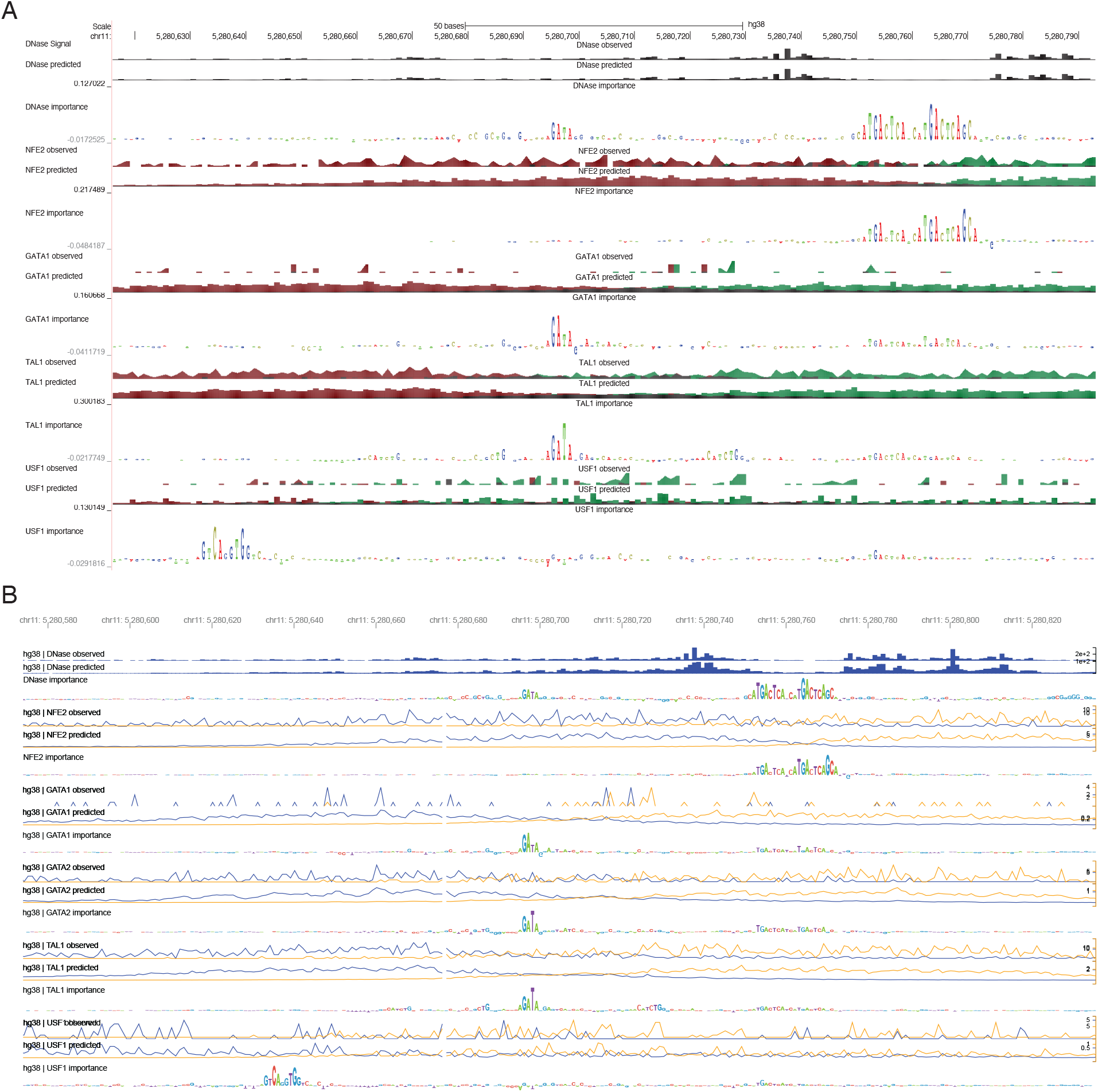
UCSC Genome Browser and Resgen session for cis-regulatory element dissection. Same tracks as in **Figure 1** in **(A)** UCSC Genome Browser and **(B)** Resgen around hg38 chr11:5280600-5280800. For ChIP-seq observed and predicted tracks, the red track denotes the plus (+) strand and the green track denotes the minus (−) strand in UCSC **(A)** and blue and orange respectively in Resgen **(B)**.

**Supplementary Table 1.** Supported functionalities of current implementations of the dynseq track.

|  | WashU | UCSC | HiGlass/Resgen |
| --- | --- | --- | --- |
| Input format | BigWig | BigWig | BigWig |
| Auto or fixed y-axis | ✓ | ✓ | ✓ |
| Reverts to BigWig display when zoomed out | ✓ | ✓ | ✓ |
| Support for files on remote servers | ✓ | ✓ | ✓ |
| Vector export | ✓ | ✓ | ✓ |
| Customizable background color | ✓ | ✗ | ✗ |
| Customizable font and nucleotide colors | ✗ | ✗ | ✓ |
| Custom fasta file per dynseq track | ✗ | ✗ | ✓ |

